# Genotypic variation in tolerance to salinity in Pakistani okra (*Abelmoschus esculentus* L.) varieties as assessed at seed germination, seedling growth and biochemical characters

**DOI:** 10.1101/2022.09.14.508060

**Authors:** Hayat Ullah, Tour Jan, Fazal Wahid, Muhammad Zahoor, Shahab Uddin, Shabana Bibi

**Affiliations:** Department of Botany, University of Malakand, Chakdara, Dir (L), Khyber Pakhtunkhwa, Pakistan; Department of Biochemistry, University of Malakand, Dir (L), Ckakdara, Pakistan

**Author notes:** (H.U.); (F.W.), (S.B.), (M.Z.).

## Abstract

Salt stress is one of the major abiotic stresses that causes reduction in crops yield. Okra (*Abelmoschus esculentus* L.) is a high-value nutraceutical vegetable because its various parts are used for different purposes. This study was conducted to calculate the salt tolerance among thirteen okra varieties. Different salt (NaCl) levels: 0, 50, 75 and 100 mM were selected to measure the response of the okra varieties to stress. The experimental data showed that all varieties were affected by salt level with a differential variation in their stress response, demonstrating the presence of genetic variability. Five varieties: “NAYAB-F1”, “Arka anamika”, “MALAV-27”, “Sarhad Green” and 051-F1 showed germination at all stress levels (0, 50, 75 and 100 mM), six varieties: “Feveeri Green”, “Punjab Selection”, “Local Multani”, “Shehzadi”, “Green Star” and “Hunza” showed germination at (0, 50 and 75 mM) and two: “Anmol” and “Sabz Pari” showed germination at (0 and 50 mM). As a result of salt stress, germination percentage (PG), leaf fresh and dry weight (LFW and LDW), shoot fresh and dry weight (SFW and SDW), root fresh and dry weight (RFW and RDW) were significantly reduced with increasing stress level. Based on the performance of variety to salt stress, five varieties were selected for biochemical analysis, concentrating on the determination of osmolytes. The values of sugar and proline were affected both by the varieties and salt levels. Variety “NAYAB-F1” showed higher sugar and proline content at all stress levels compared to varieties “Arka anamika”, “MALAV-27”, “051-F1” and Sarhad Green. Correspondingly, in the stressed seedling a decreasing trend in chlorophyll “a” and “b” were noted depending on the varieties and stress concentrations. After a series of experiment, it has been concluded that varieties “NAYAB-F1”, “Arka anamika”, “MALAV-27”, “Sarhad Green” and “051-F1” were recommended as salt tolerant varieties and could be utilized in the breeding program of salt tolerant okra.

## 1. Introduction

Soil salinization is one of the most important abiotic stresses that undesirably upset plant growth and development around the world [1, 2]. It has been described that nearly 19.5% of all irrigated land and 2.1% of dry land is affected by salt stress [3]. NaCl is the common salt at current and it has always been the focus of salinity research [4, 5]. Surplus salt concentration is toxic inside plant cells due to ionic stress and smashes the equilibrium of ions [6]. Soil salinization increases the osmotic pressure, as consequence the plant root cannot absorb water [7]. Salt stress inhibited Carbon fixation as excessive salt level might result in the closing of stomata which finally decreases the carbon dioxide availability in the leaves) [8]. Salinity stress have numerous damaging effects on plants which comprises root inadequate growth, flowering inhibition, browning or burning of leaf tip or margins, reduced vigor, and reduction of crops productivity [9]. Salinity stress shrink water opportunity and generates toxic ions such as Na^+^ and Cl^-1^ around the root area, as result the growth of plant was reduced [10]. Seed germination is one of the most essential and vital phases in the growth cycle of a plant that determines the yield as germination is the first stage of a plant’s life cycle for establishment of a crop [11, 12]. Successful seed germination and strong seedling potency are critical determinants for crop growth in their success, as these parameters lead to consistent plant growth and, consequently, high production [13]. Saline soil has opposing effect on germination rate, seedling growth, fresh and dry matter of seedlings, seed viability, and chloroplast pigments as well as chlorophyll and carotenoid biosynthesis [14]. It has been reported that salt stress delayed seed germination of most crop i.e. melon [15] and tomato. Soil salinity is one of the major physical factors of an environment, which determines the success and failure of plants establishment. Many research studies have documented the plant defensive replies to salt, important to the establishment of plants with improved salinity tolerance [16, 17]. Soil salinization interrupts the root’s functioning for minerals from soil to plant transpiration stream with dangerous damage, [18] and disrupts ionic homeostasis [19]. Moreover, this will result in the increased photorespiration and reduced photosynthesis, deteriorated chlorophyll synthesis, [20, 21]. In reply to environmental stresses, plants produced some defensive compounds. The assemblage of osmo-regulatory compounds and specific ions absorption are required for defensive responses in plants against ions surplus toxicity and osmotic stress [22]. Plants in saline soil frequently meet osmotic stress and ion toxicity. Still plants have many defense mechanisms to adapt these environments, [23] and assemblage of osmo-protectant compounds such as proline, soluble sugars, and glycine betaine to continue balanced water uptake in plants [24, 25].

Many authors have been reported that salinity badly damage the process of germination in several plants such as *Oryza sativa* [26], *Triticum aestivum* [27], *Zea mays* [28, 29] and *Brassica* spp. [30]. Okra (*Abelmoschus esculentus* Moench) is one of the most important nutritious vegetable crops around the world [31]. It is a ridiculous source of nutrients like Calcium, potassium vitamins, carbohydrates, and unsaturated fatty acids, [32]. The reply of okra crop to salt stress is variable, from very sensitive to moderately sensitive [33]. Keeping in view the objective that the increase of salt resistant varieties is one of the real source for the increasing of okra yield on saline soils. In this investigation thirteen okra varieties were exposed to salt (NaCl) stress at changing levels (0, 50, 75, 100 mM), and their reply to stress was systematically evaluated on the basis of characters concerned to seed germination, seedling growth and synthesis of biochemical compounds, namely sugar, proline and chlorophyll “a” and “b” that are usually employed as indicators of salt tolerance.

## 2. Materials and Methods

### 2.1. Collection of experimental materials

Healthy seeds of thirteen (13) different okra (*Abalmoschus esculentus*) varieties were obtained from local market, Battkhela, Khyber Pakhtunkhwa, Pakistan (Table 1).

**Table 1.** List of varieties used in the experiment

| S. No | Variety Names | S.No | Variety Names |
| --- | --- | --- | --- |
| 1 | NAYAB-F1 | 8 | Sabz Pari |
| 2 | Arka anamika | 9 | Feveeri Green |
| 3 | NALAV-27 | 10 | Shehzadi |
| 4 | Punjab selection | 11 | Hunza |
| 5 | Local Multani | 12 | Green star |
| 6 | Sarhad green | 13 | Anmol |
| 7 | 051-F1 |  |  |

### 2.2. Pots experiments

Experiments were conducted in plastic pots (depth 23 cm, diameter 19 cm) each containing of 5 kg of garden soil. Five okra seeds were placed in each pot and irrigated with different solutions of salt or tap water (control). The experiment was conducted to determine the effects of salt stress on germination indexes as well as on morphological and physiological characters of *Abelmoschus esculentus* L. In the current experiment (0, 50, 75, and 100 mM NaCl) concentrations were used to screen out salt tolerant varieties. These experiments were conducted at the month of April and May respectively. The experiment was conducted in randomized complete block design having three replicates. After 72 hours of sowing, seed germination was recorded randomly. After 15 days of germination morphological and physiological characters was determine.

### 2.3. Germination assessment

The number of germinated seed was recorded daily up to ten days after 72 hours of sowing. From germination count, germination responses were recorded to screen out salt tolerant, moderat tolerant and sensitive varieties, based on germination percentage and germination index (GI) [34].

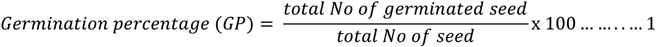

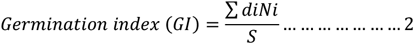

Where, di= Number of days after sowing seeds under a particular treatment, Ni= the number of germinated seeds and S= the total number of seeds sown for the experiment.

### 2.4. Germination energy (GE)

Germination energy was calculated with the help of following formula [35],

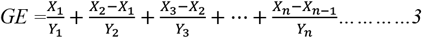

Hence, X_n_ is the numbers of seed germination on last (nth) counting date and Y_n_ represents the number of days from sowing to last (nth) counting date.

### 2.5. Vigor index (VI)

For the determination of vigor index (VI), first we calculated seed germination and then measure root and shoot length in centimeter. For the calculation of seed vigor index the following formula was used,

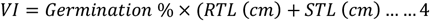

Where, ‘VI’ is the vigor index ‘RTL’ is the root length and ‘STL’ is the shoot length.

### 2.6. Morphological responses

The morphological characters i. e. fresh and dry weight of leaf, shoot and root respectively were recorded.

### 2.7. Assessment of salts stress tolerance based on biochemical responses

Based on the performance of thirteen okra varieties to salt stress, five varieties were selectd for biochemical evaluation. Variety “NAYAB-F1”, “Arka anamika”, “051-F1”, “Sarhad Green” and “MALAV-27” were evaluated, using the seedling tisues in sugar, proline and chlorophyll “a” and “b” evaluation.

#### 2.7.1. Sugar content

Sugar content of fresh leaf was calculated by the method of [36]. Fresh leaf weight 0.2 g was heated with 10 mL ethanol (80%) in a water bath at 95C° for 10 minutes. Then 1 mL of phenol (18%) was added to 0.5 mL of the prepared sample extract and then it was incubated at room temperature for 1 hour. After that 2.5 mL of H_2_SO_4_ was added to it, and then vortex and the absorbance of the supernatant was read at 490 nm on UV Spectrophotometer.

#### 2.7.2. Proline Content

Proline content of leaf was determined by the method of [37]. Fresh leaves (0.2 g) was weighted and put in 10 ml 3% sulfosalicylic acid solution in test tubes for 48 hours to adjust the supernatant. Then 2 ml supernatant was taken from prepare sample in test tube, and then 2 ml of glacial acetic acid, 2 ml of ninhydrine reagent was added with this supernatant to make the solution. The solution was shacked and heated at 100 °C in water bath for one hour. After heating the solution was cooled and 4 ml toluene was added to each test tube. Toluene was used as a blank sample. Absorbance of supernatant was measured at 520 nm on UV spectrophotometer.

#### 2.7.3. Chlorophyll Content

Chlorophyll content of leaves was determined by the method of [38]. Fresh leaves weight 0.2 g was grinded and then dipped in ethanol (80%) in test tube. The test tube was heated in water bath at 80°C for 15 minutes. After that the test tube was centrifuge at 10,000 rpm for 6 minutes and the absorption of the supernatant was noticed through a spectrophotometer.

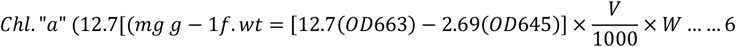

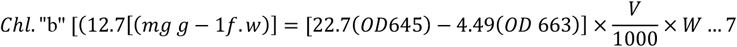

Where V = Volume of the extract (mL) and W = Weight of the fresh leaf tissue (g).

### 2.8. Data analysis

The mean data from three replications were analyzed by the method of one-way analysis of variance (ANOVA) using SAS statistical Software. Varieties were ranked according the germination percentage over different salt concentration in distal water across one trail month.

## 3. Results

### 3.1. Impact of salt levels on characters associated to seed germination and seedling growth

To calculate the reaction of Pakistani okra varieties (Table1) to salt stress, seeds were germinated on different salt levels (0, 50, 75 and 100 mM), then seed germination and seedling growth potential were evaluated. The experimental data revealed a significant result of the salt stress level on seed germination and seedling growth potential of the tested okra varieties, with the stress effects being in maximum cases proportional to the stress level applied, leading to severe effects at high level for all the characters under study (Table 2 and 3). The stress level greatly affected seed GP, demonstrating decreasing tendencies as salt level increased (GP: 80.02, 41.03, 24.10 and 3.68% at 0, 50, 75 and 100 mM respectively) (Table 2). Severe effects of stress were also observed in growth index (GI), which has been greatly affected by the stress level (Table 2). GI showed decreasing affinities as salt level increased, ranged from 4.31 to 0.46% compared to the control, hence showing the disparity effects of different stress level on germination (Table 2). Significant effect of the stress level on the germination energy (GE) and vigorous index (VI) were also observed, with the effects being proportional to the stress level used. The GE and VI dropped from 1.88 to 0.23% and 574.21 to 16.89% as salt level increased from 0 to 100 mM (Table 2). The effect of salt stress was also evaluated on the seedling growth and the effect was observed proportional to the stress level applied. The leaf fresh and dry weight (LFW & LDW), root fresh and dry weight (RFW & RDW) decreased as salt stress level increased and thus the differential effects of salt stress level on seedling growth was observed (Table 2). Hence, LFW and LDW decrease with increasing salt level. Significant differences between the salt stress levels (0. 50, 75 and 100 Mm) were observed, where in the condition of SDW and RDW, no significant differences were observed amongst the stress levels (0, 50, 75 and 100 mM).

**Table 2.**
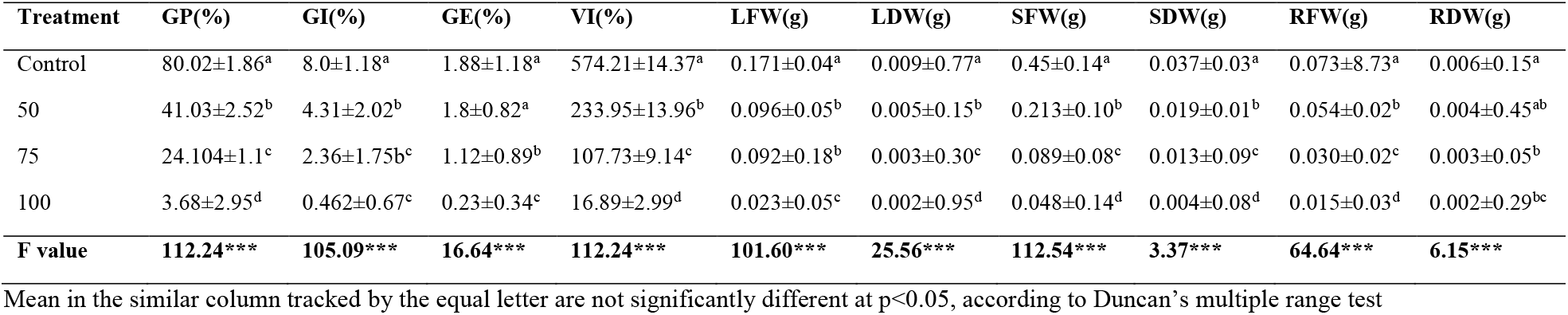
Effect of the salt levels on seed germination and seedling growth, regardless of okra varieties

**Table 3.**
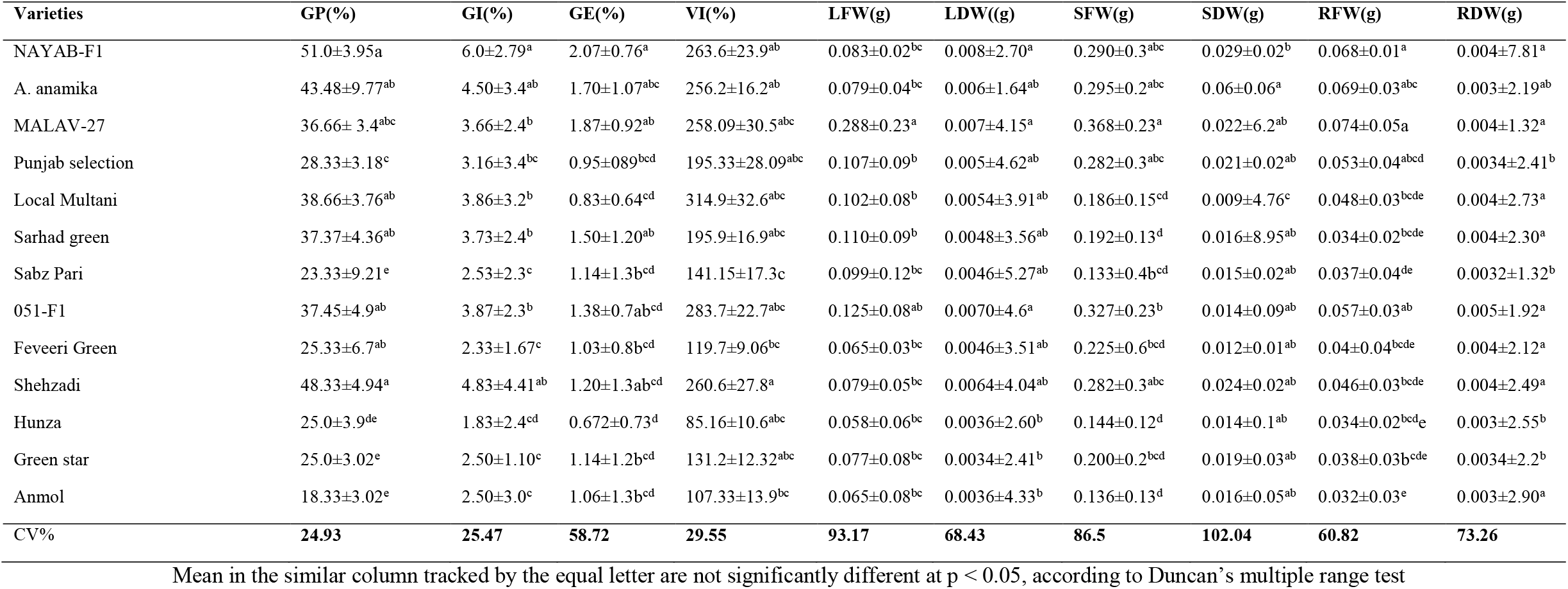
Effect of the okra varieties on seed germination and seedling growth, regardless of the salt stress.

### 3.2. Impact of okra varieties on characters associated to seed germination and seedling *growth*

The experimental data show the substantial effect of okra variety on seed germination and seedling growth under salt stress condition (Tables-2 and-3). The inconsistent replies of the varieties to salt stress was demonstrated for all the characters under study (Table 3). GP and VI was considerably affected by the variety, as showed by the mean GP (Table 3), which are ranged from 51.0% to 18.33% in “NAYAB-F1” and “Anmol” varieties, respectively. Such values indorsed a salt tolerance capability for “NAYAB-F1” and sensitivity for “Anmol”, they are endorsed to their inherent germination potential, as showed by the differences in GP in the corresponding control values, for “NAYAB-F1” 93.33% and “Anmol” 86.6% (Table 4). The other varieties “Shehzadi”, “Arka anamika”, Local Multani”, “051-F1”, “Sarhad Green”, “MALAV-27” and “Punjab Selection” were characterized by GP ranging from 48.33% to 28.33, while the other varieties “Sabz Pari”, “Green Star”, “Anmol” and “Feveeri Green” exhibited intermediate GP that ranging from 25.23 % to 18.33% (Table 3). Comparison of the seedling growth showed that the salt stress effect was more severe on RDW than on SDW (Table 3). While the stress effects were obvious in the studied varieties. So, differential effects were observed for SFW and RFW, with “MALAV-27” displaying the maximum value for both characters, while the minimum values were recorded in varieties “Sabz Pari” and “Anmol, respectively (Table 3). The data highlighted in table-3 show a differential variation for the LDW, SDW and RDW. In relation to SDW, “A. anamika” followed by “NAYAB-F1”, “MALAV-27”, “051-F1” and “Shehzadi” showed the highest tolerance to salt stress, however “Feveeri Green” ranked as the low performing variety. In contrast, “051-F1”, “Shehzadi” “MALAV-27” “Sarhad Green” and “Local Multani” were proved to be the salt sensitive varieties based on LDW and RDW values (Table-3).

**Table 4.**
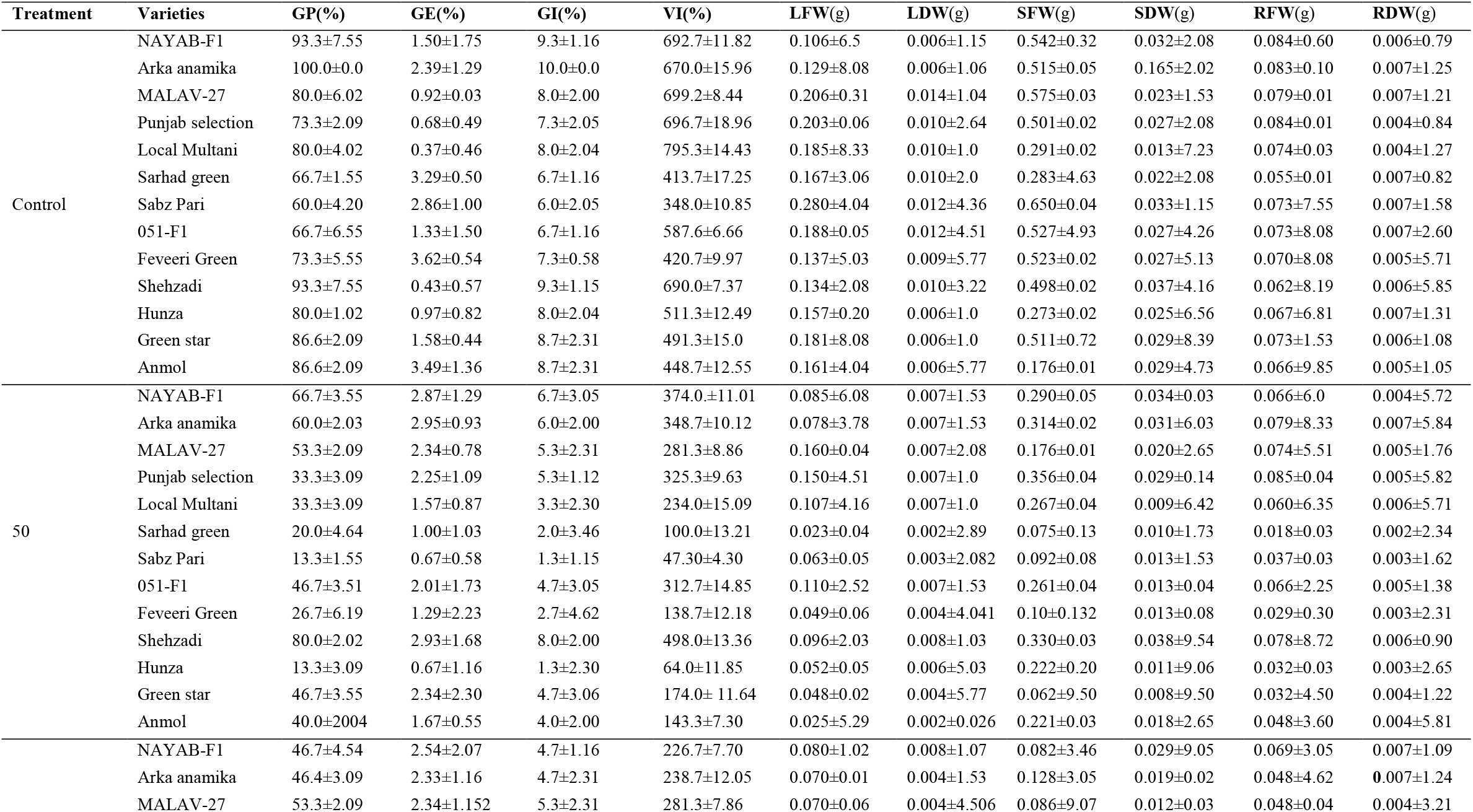
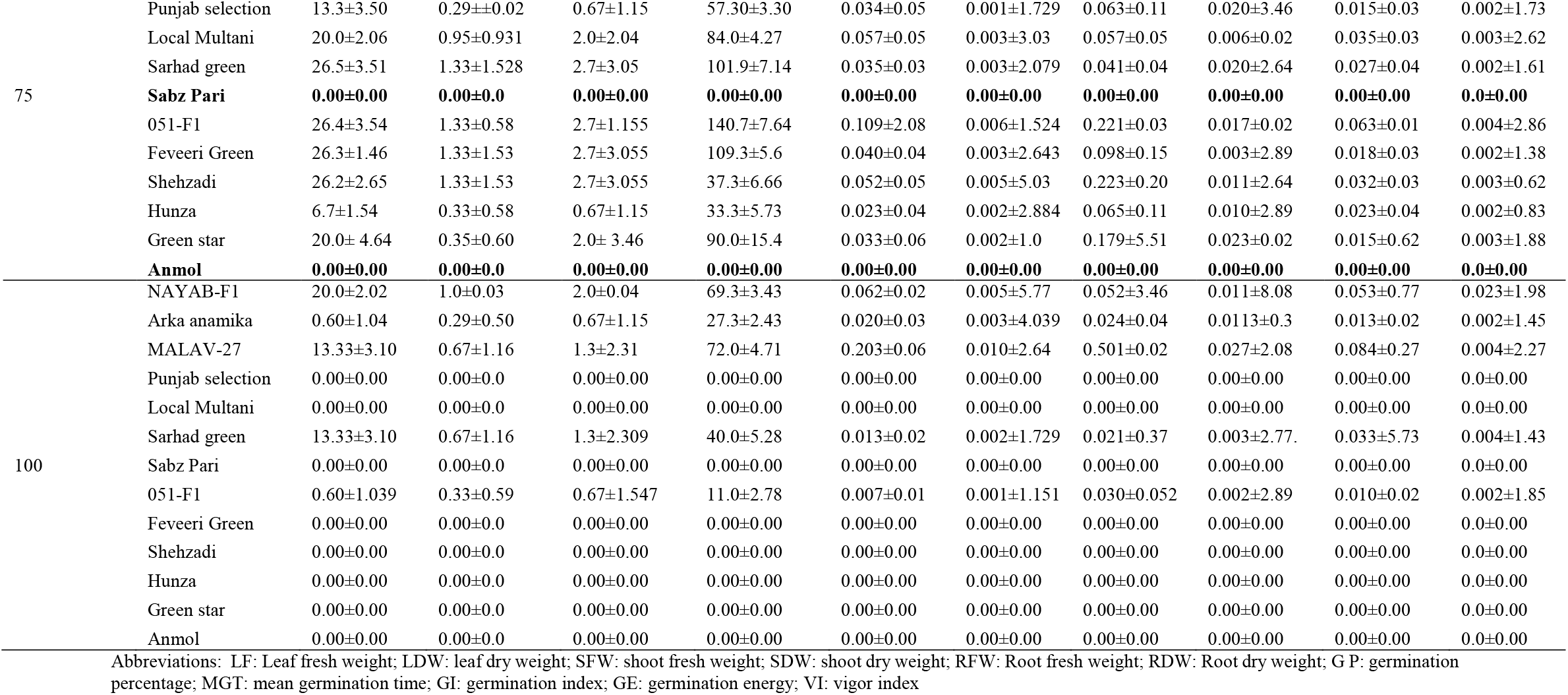
The mean effect of okra varieties on seed germination and seedling growth, regardless of different level of salinity.

### 3.3. Influence of the varieties and salt stress levels on character related to the germination of seed and seedling growth

The investigation of variance applied on individual data, all the characters related to seed germination and seedling growth in stress environment were affected by the variety and salt levels (Table-4). In the lack of salt stress, germination was significantly affected by the varieties, thus certifying an inconstant germination potential per seed, which could be attributed to the average strength of seeds. Among these thirteen varieties under normal condition, varieties “Arka anamika” and “Sabz Pari” showed the maximum and minimum GP (100.0% and 60.00 %, correspondingly) (Table 4). With salt stress, the germination of all okra varieties was greatly disturbed and the effects were equivalent to its levels. At the level of 50 mM, “Shehzadi”, “NAYAB-F1”, “Arka anamika” and “MALAV-27” showed the highest GP values ranging from 80.00 % to 53.3%, while “051-F1”, “Green Star”, “Anmol”, “Punjab Selection” and “Local Mulatani” showed moderate GP value ranged from 46.7 to 33.3% and “Feveeri Green”, “Sarhad Green”, “Sabz Pari” and “Hunza”, showed the lowest GP values ranging from 20.0% to 13.33% respectively. At the stress level of 75 mM, variety “MALAV-27”, “NAYAB-F1”, and “Arka anamika” showed the highest GP values ranged from 53.33% to 46.47%, variety “Sarhad Green”, “051-F1”, “Feveeri Green”, and “Shehzadi”, presented moderate GP values ranged from 20.0% to 26.67%), variety “Local Multani”, “Green Stare” showed lowest GP value 20.00%, and variety “Sabz Pari” and “Anmol” showed no germination (table 4). At highest stress level 100 mM, only “NAYAB-F1”, “Sarhad Green” and “MALAV-27” showed 20.0% and 13.33% germination respectively (Table 4). Similarly, the response of maximum varieties to salt stress showed a reduction in SFW and RFW (Table 4). The SFW and RFW presented a decreasing trend, as salt level increased. The highest values for these characters were recorded in “MALAV-27” and “Arka anamika” whereas the lowest values were noted in “Sarhad Green” and “Anmol” (Table 4).

### 3.4. Effect of the salt stress level on the content of osmolytes and chlorophyll “a” and “b”

The data in table-5 highlighted that the various levels of salt stress affected the content of sugar, proline and chlorophyll “a” and “b” of okra varieties. The sugar content, at the level of 50 and 75 mM, showed a differential effect among thirteen varieties were manifested in stressed seedlings as compared to the control ranging from 0.370 to 382 μg g-^1^ FW. The maximum sugar content (0.382 μg g-^1^ FW) was originated at stress level 75 mM, while at 50 mM stress level the respective values of sugar content was 0.3700 μg g-^1^ FW, however the content of sugar was minimum (0.147 ug g-^1^ FW) at 100 mM stress level. The content of proline was maximum (0.339 μmol g-^1^ FW) at 50 mM stress level, whereas the respective values of proline at 75 mM stress level was 0.338 μmol g-^1^ FW (Table 5). The minimum value of proline content 0.132 μmol g-1 FW was noted at stress level of 100 mM. As expected, significant reduction in chlorophyll “a” and “b” was noted in stressed seedling, as compared to the control (Table-5). The value of chlorophyll “a” ranged from 15.49 to 1.22 mg g-1 FW, while chlorophyll “b” value ranged from (14.66 to 1.46 mg g-1 FW) at (0 to 100 NaCl level) (table 5).

**Table 5.** Effect of salt levels on the content of osmolytes and chlorophyll (“a” and “b”), irrespective of the okra varieties

| Salt level (mM) | Sugar $\mu\text{g g}^{-1}$ FW | Proline $\mu\text{mol g}^{-1}$ FW | Chl. “a” $\text{mg g}^{-1}$ FW | Chl. “b” $\text{mg g}^{-1}$ FW |
| --- | --- | --- | --- | --- |
| Control | 0.350±0.10<sup>b</sup> | 0.266±0.04<sup>b</sup> | 15.49±0.22<sup>a</sup> | 14.66±0.70<sup>a</sup> |
| 50 | 0.3700.22<sup>a</sup> | 0.339±0.19<sup>a</sup> | 7.032±1.79<sup>b</sup> | 9.18±0.95<sup>b</sup> |
| 75 | 0.382±0.25<sup>a</sup> | 0.338±0.23<sup>a</sup> | 4.69±0.83<sup>c</sup> | 5.87±1.06<sup>c</sup> |
| 100 | 0.147±0.32<sup>c</sup> | 0.132±0.29<sup>c</sup> | 1.22±0.68<sup>d</sup> | 1.46±1.29<sup>d</sup> |
| **F Value** | **58.39\*\*\*** | **20.75\*\*\*** | **143.12\*\*\*** | **112.42\*\*\*** |

### 3.5. Effect of variety on the content of osmolytes and chlorophyll “a” and “b”

Results of the physiological data in table-6 show a significant effect of the variety on the content of sugar, proline and chlorophyll “a” and “b” in okra seedlings (Table 6). The highest sugar content was noted in three varieties, in “NAYAB-F1” (0.590 μg g-1 FW), “MALAV-27” (0.498 μg g-1 FW) and “Arka anamika” (0.469 μg g-1 FW), while lowest sugar content was noted in two varieties, in “Sabz Pari” (0.138 μg g-1 FW) and “Anmol” (0.164 μg g-1 FW). However, in rest of the varieties, the sugar content was not significantly different. The variety “NAYAB-F1” was further recognized by the highest value of proline content (0.452 μmol g-^1^ FW). On the other hand, variety “Sabz pari” showed the lowest content of proline (0.161μmol g-^1^ FW) (Table 6). In relation with the content of chlorophyll “a” and “b”, the highest values were noted in “NAYAB-F1” (11.47 and 11.04 mg g-1 FW) and in “051-F1” (11.36 and 11.69 mg g-1 FW respectively), whereas the lowest values were noted in “Anmol” (4.42 and 4.10 mg g-1 FW, respectively) (Table 6).

**Table 6:** Effect of okra varieties on the content of osmolytes and chlorophyll “a” and “b” irrespective of the salt levels

| Varieties | Sugar $\mu\text{g g}^{-1}$ FW | Proline $\mu\text{mol g}^{-1}$ FW | Chl “a” $\text{mg g}^{-1}$ FW | Chl “b” $\text{mg g}^{-1}$ FW |
| --- | --- | --- | --- | --- |
| NAYAB-F1 | 0.59±0.14<sup>a</sup> | 0.46±0.11<sup>a</sup> | 11.47±0.95<sup>a</sup> | 11.04±0.14<sup>ab</sup> |
| Arka anamika | 0.47±0.12<sup>ab</sup> | 0.39±0.07<sup>ab</sup> | 10.82±2.30<sup>b</sup> | 9.79±2.53<sup>b</sup> |
| MALAV-27 | 0.49±0.14<sup>ab</sup> | 0.33±0.06<sup>abcd</sup> | 8.41±2.84<sup>ab</sup> | 8.77±1.52<sup>ab</sup> |
| Punjab selection | 0.32±0.19<sup>bcd</sup> | 0.23±0.16<sup>bcd</sup> | 5.08±1.69<sup>d</sup> | 5.35±0.59<sup>cdef</sup> |
| Local Multani | 0.30±0.17<sup>abc</sup> | 0.29±0.17<sup>ab</sup> | 7.66±2.73<sup>bcd</sup> | 8.02±1.47<sup>c</sup> |
| Sarhad green | 0.37±0.05<sup>bcd</sup> | 0.33±0.06<sup>ab</sup> | 8.94±6.11<sup>ab</sup> | 8.55±1.05<sup>ab</sup> |
| Sabz Pari | 0.13±0.13<sup>d</sup> | 0.16±0.15<sup>d</sup> | 7.04±7.28<sup>bcd</sup> | 6.66±2.94<sup>bcd</sup> |
| 051-F1 | 0.37±0.09<sup>bcd</sup> | 0.34±0.09<sup>abc</sup> | 11.36±2.29<sup>a</sup> | 11.69±0.96<sup>a</sup> |
| Feveeri Green | 0.24±0.16<sup>cd</sup> | 0.21±0.16<sup>bcd</sup> | 6.77±2.10<sup>c</sup> | 5.97±1.82<sup>cd</sup> |
| Shehzadi | 0.28±0.19<sup>bcd</sup> | 0.28±1.9<sup>abcd</sup> | 7.19±1.94<sup>bcd</sup> | 7.33±0.95<sup>bcd</sup> |
| Hunza | 0.15±0.10<sup>d</sup> | 0.20±0.16<sup>abcd</sup> | 5.33±2.50<sup>cd</sup> | 4.55±1.22<sup>ef</sup> |
| Green star | 0.21±0.17<sup>cd</sup> | 0.22±0.18<sup>bcd</sup> | 5.86±1.05<sup>cd</sup> | 6.40±2.09<sup>ab</sup> |
| Anmol | 0.16±0.19<sup>cd</sup> | 0.16±1.95<sup>cd</sup> | 4.42±1.41<sup>d</sup> | 4.10±0.96<sup>ef</sup> |

### 3.6. Effect of variety and salt stress level on the content of osmolytes and chlorophyll “a” and “b”

Data in table 7 showed that the contents of sugar, proline and chlorophyll “a” and “b” were affected by the variety and salt stress level applied. The sugar concentration was found higher in variety “NAYAB-F1” compared to varieties Arka anamika”, “MALAV-27”, “051-F1” and Sarhad green. The variety “NAYAB-F1” showed a gradual rise in sugar content as stress level increased. The concentration of sugar in varieties “Arka anamika”, “MALAV-27”, “051-F1” and “Sarhad green” increased up to 75 mM NaCl concentration, thereafter, a decline was observed at 100 mM NaCl concentration compared to the control values (Table 7). Regarding proline, variety “NAYAB-F1”, presented the highest proline content, at all stress levels, variety “Arka anamika”, “MALAV-27”, “051-F1” and “Sarhad green” showed highest proline content at 50 and 75 mM but at 100 mM stress level lowest value was presented as compared to the control (Table-7). Correspondingly, the values of chlorophyll “a” and “b” was differentially affected both by the varieties and salt levels. Chlorophyll “a” and “b” was reduced in the stressed seedling as compared to the control, yet a decreasing trend were noted depending on the variety and stress amount (Table 7). Specifically, “NAYAB-F1” presented the highest values of chlorophyll “a” and “b” across all levels of salt, among all varieties individually. While “Sarhad green” showed the lowest values of chlorophyll “a” and “b” at all stress levels (Table 7).

**Table 7.** Responses of okra varieties to salt stress in relation to the content of sugar, proline and chlorophyll “a” and “b”

| Varieties | Salt levels | Sugar $\mu\text{g/g}$ | Proline $\mu\text{mol/g}$ | Chl “a” $\text{mg/g}$ | Chl “b” $\text{mg/g}$ |
| --- | --- | --- | --- | --- | --- |
| NAYAB-F1 | Control | 0.36±0.02 | 0.30±1.13 | 15.66±0.59 | 16.78±2.06 |
| | 50 | 0.54±0.01 | 0.48±2.51 | 12.30±0.91 | 13.57±0.54 |
| | 75 | 0.63±0.01 | 0.47±1.01 | 9.80±0.18 | 11.42±0.57 |
| | 100 | 0.74±0.02 | 0.51±0.01 | 7.97±0.26 | 7.56±1.49 |
| Arka anamika | Control | 0.37±0.08 | 0.32±0.85 | 17.91±1.36 | 16.41±0.60 |
| | 50 | 0.52±0.46 | 0.40±0.02 | 11.13±0.96 | 12.38±1.86 |
| | 75 | 0.68±0.64 | 0.45±0.05 | 7.69±0.75 | 10.60±1.30 |
| | 100 | 0.48±0.53 | 0.28±0.49 | 2.252±1.49 | 2.81±1.14 |
| MALAV-27 | Control | 0.46±0.02 | 0.21±0.24 | 16.93±0.91 | 15.72±0.13 |
| | 50 | 0.60±0.02 | 0.39±0.01 | 10.89±1.22 | 11.32±0.79 |
| | 75 | 0.62±0.52 | 0.32±0.28 | 5.83±1.89 | 5.97±0.381 |
| | 100 | 0.40±0.53 | 0.26±0.43 | 2.60±1.81 | 3.18±2.50 |
| Sarhad Green | Control | 0.35±1.50 | 0.28±0.61 | 15.13±0.28 | 14.84±0.91 |
| | 50 | 0.36±0.31 | 0.28±0.24 | 6.29±1.90 | 5.637±2.76 |
| | 75 | 0.38±0.33 | 0.423±0.36 | 5.50±0.63 | 4.457±1.94 |
| | 100 | 0.32±0.51 | 0.19±0.50 | 2.14±0.72 | 2.55±0.41 |
| 051-F1 | Control | 0.33±0.03 | 0.27±0.21 | 16.97±0.16 | 15.75±1.78 |
| | 50 | 0.40±0.01 | 0.42±9.16 | 11.02±0.38 | 12.43±0.56 |
| | 75 | 0.51±0.01 | 0.48±0.61 | 10.05±0.87 | 10.97±0.36 |
| | 100 | 0.39±0.44 | 0.22±0.46 | 3.06±1.29 | 3.94±1.81 |

### 3.7. Analysis of variance

The ANOVA of germination indexes show that treatment, varieties and the interaction between treatment and varieties was less significant in GE and VI, while the interaction between treatment and varieties in GI, GP was non-significant. Similarly, analysis of variance regardless morphological characters shows that treatment, varieties and the interaction between treatment and varieties in LFW and LDW), SHFW and SHDW, RFW and RDW, were highly significant (Table 8).

**Table 8.** Analysis of variance for characters related to agronomical and morphological parameters under stress condition

| S.O.V. | DF | GP | GI | GE | VI | LFW | LDW | SFW | SDW | RFW | RDW |
| --- | --- | --- | --- | --- | --- | --- | --- | --- | --- | --- | --- |
| Varieties | 13 | 32303.5*** | 80.76*** | 80.76*** | 12.27*** | 4.56*** | 9.013*** | 0.093*** | 0.072*** | 6.94*** | 0.002*** |
| Treatment | 4 | 1954.4*** | 4.87*** | 2.164* | 6639*** | 0.164*** | 4.665*** | 1.299*** | 0.0053* | 0.027*** | 2.146*** |
| Treatment *Varieties | 52 | 157.75*** | 1.357<sup>NS</sup> | 2.006*** | 1175* | 0.003*** | 2.460* | 0.028*** | 0.0018<sup>NS</sup> | 0.0007*** | 7.612*** |
| **CV%** |  | **52.2** | **52.2** | **78.12** | **50.36** | **32.82** | **71.15** | **26.1** | **185.2** | **35.9** | **88.9** |

S.O.V.: source of variance; DF: degree of freedom; CV: coefficient of variance; GP: germination percentage; GI: germination index; GE: germination energy; VI: vigor index; LFW leaf fresh weight; LDW: leaf dry weight; SFW: shoot fresh weight; SDW: shoot dry weight; RFW: root fresh weight; RDW: root dry weight.

## 4. Discussion

Salinity poses severe limitations to seed germination and seedling growth and the selection of a salt resistant variety is great significant in breeding programs when expecting the development of salt resistance varieties. In this investigation we have tested three levels of salt at the stage of germination and early seedling growth to identify the level of salt tolerance in thirteen okra varieties and to select the resistance varieties by measuring specific characters associated with seed germination, seedling growth and determination of biochemical compounds (sugar, proline and chlorophyll “a” and “b”) that have an important role in the ability to resist stress. At the stage of germination, the experiment exposed that salt stress induced lower GP, GI, GE and VI of okra varieties. In relation with germination characters, various stress levels of tolerance were evaluated among the varieties under study, variety “NAYAB-F1” verifying to be the tolerant. The varieties which are minimum affected by salt stress may be probable sources of gene for salt tolerance, such varieties are significant in plant breeding programs as confirmed in tomato [39]. So, these varieties which germinate at a high salt stress may be used as potential contributors for salinity tolerance. In general, the data presented in (table 2 and 3), there is a disparity in the studied characters associated with seed germination and seedling growth of okra varieties at any level of stress, the overall trend is that salt stress differentially affects all studied characters, with the effects being in maximum cases proportional to the stress level applied. These results are dependable with earlier studies regarding the effects of salt stress on the seed germination of lettuce [40], rice [41], chickpea [42] and wheat [43]. Similar results of germination inhibition were also highlighted by [44–46] in other plants. Our results indicate that a balance increasing severity of effects depending on salt concentration, thus proving the earlier results that the variable stress levels differentially affect germination and seedling growth in different plants species, sugar beet and cabbage [47], soybean [40] and lentil [48]. All the studied varieties in our investigation were badly affected by salt stress, particularly at stress level 100 mM NaCl.

Salinity stress was clearly demonstrated on early seedling growth, obvious in terms of the reduction in the fresh and dry weight of leaf, root and shoot of the studied varieties. Our results in (Table 3) indicating that increasing salt stress level caused a decreasing tendency in seedling fresh and dry weight of root and shoot, which could be credited to ionic effects occurring as a result of a proportional increase in Na^+^ level [49]. It was also reported that the inhibition of a plant shoot and root growth is one of the key agricultural indicators to salt stress [50]. Our results demonstrated that high stress level 100 mM drastically affected seedling growth. However, the rate of effect varied among the different varieties. This disparity indicates the different levels of salt tolerance in the studied varieties. In accord with our result, other studies on salt stress indicated biomass reduction as salinity increases [51, 52].

Proline is an important osmolytes that accumulate in plants under several environmental stresses, including salinity [53, 54]. The present investigation showed that varieties “051-F1”, “Arka anamika”, “NAYAB-F1” and “MALAV-27” showed the high proline content in leaves under salinity stress than those of “Green star”, “Feveeri green” and “Hunza” varieties.

Sugar is a significant osmotic regulator used by plants to resist stress condition, and their assemblage in plants facing salinity stress is an adaptive mechanism [5, 54]. Indeed, in our results, salt stress produced increase sugar content as compared to the control plants. Again varieties “NAYAB-F1”, “MALAV-27”, “Arka anamika” and “051-F1” showed highest sugar content.

In the process of photosynthesis, the green pigment chlorophyll “a” and “b” act as photoreceptors in plant, play an important role and absorbs the light [55]. In our study with salt stress the varieties namely NAYAB-F1, 051-F1, “Arka anamika”, “MALAV-27” and “Sarhad Green” showed higher chlorophyll a and b content, while varieties namely “Local Multani”, “Shehzadi”, “Sabz Pari” and “Feveeri Green” showed moderate chlorophyll “a” and “b” content, and varieties namely “Green Star”, “Hunz” and “Anmol” showed lower chlorophyll “a” and “b” contents. It has been reported in the literature that salt stress tolerant genotypes showed greater chlorophyll contents as compared to salt stress susceptible lines [56, 57]. Our data showed a general decreasing trend of chlorophyll “a” and “b” in stressed plants, as compared to the control environment, which indicates that the response to high salinity may involve a decrease in chlorophyll content in various plant species, including pepper [58].

## 5. Conclusion

According to the result of this experiment, it can be concluded that reduction in germination percentage, seedling fresh and dry weight of leaf, root and shoot was strongly connected with salt stress. These evaluated characters can serve as an efficient tool for the calculation of salt tolerance potential for okra varieties. Definitely, the conclusion of this study can be confirmed by the results of recent investigations where we noticed that germination related traits, morphological and physiological function of okra has been decrease with increasing application of salinity level.

## Acknowledgement

The authors acknowledge the Department of Botany, University of Malakand, Chakdara Dir (L), KP, Pakistan for providing Laboratory facilities to effectively complete this work.

## Author Contributions

Conceptualization; T.J., H.U. and M.Z. Write up of original draft of paper; H.U., F.W., T.J., and M.Z. Revision of the paper; T.J., H.U. and M.Z. Methodology; H.U., T.J. and S.B. Formal analysis; H.U. and S.B. All authors have read and agreed to the published version of the manuscript

## References

1. Hafsi C, Romero-Puertas MC, Gupta DK, Luis A, Sandalio LM, Abdelly C. Moderate salinity enhances the antioxidative response in the halophyte Hordeum maritimum L. under potassium deficiency. Environmental and Experimental Botany. 2010; 69: 129–136.

2. Parida AK, Das AB, Salt tolerance and salinity effects on plants: a review. Ecotoxicology and Environmental Safety. 2005; 60: 324–349.

3. Mashali A. FAO global network on integrated soil management for sustainable use of salt affected soils. in Proceedings of International Symposium on Sustainable Management of Salt Affected Soils in the Arid Ecosystem. 1997.

4. Tavakkoli EP, Rengasamy, McDonald GK. High concentrations of Na+ and Cl–ions in soil solution have simultaneous detrimental effects on growth of faba bean under salinity stress. J. Exp. Bot. 2010; 61: 4449–4459.

5. Munns R, Tester M. Mechanisms of salinity tolerance. Annu. Rev. Plant Biol. 2008; 59: 651.

6. Tahira A, Pervez MA, Ayyub CM, Shaheen MR, Sana T, Shahid M.A, Bilal R.M, Abdul M. Evaluation of different okra genotypes for salt tolerance. International Journal of Plant, Animal and Environmental Sciences. 2014; 43: 23–30.

7. Shahid MA, Pervez MA, Balal RM, Ahmad R, Ayyub CM, Abbas T; Akhtar N. Salt stress effects on some morphological and physiological characteristics of okra (Abelmoschus esculentus L.). Soil. Enviro. 2011; 30–1.

8. Roohi A, Nazish B, Maleeha M, Waseem SA. A critical review on halophytes: salt tolerant plants. J. Med. Plant Res. 2011; 533: 7108–7118.

9. Sonon L, Saha U, Kissel D. Soil salinity testing, data interpretation and recommendations: University of Georgia Cooperative Extension Circular. 2015; 10: 6–19.

10. Silveira JAG, Araújo SA, Lima JP, Viégas RA. Roots and leaves display contrasting osmotic adjustment mechanisms in response to NaCl-salinity in Atriplex nummularia. Environ. Exp. Bot. 2009; 66: 1–8.

11. Almansouri MJ, Kinet M, Lutts S. Effect of salt and osmotic stresses on germination in durum wheat (Triticum durum Desf.). Plant Soil Environ. 2001; 231: 243–254.

12. Zhu G, An L, Jiao X, Chen X, Zhou G, McLaughlin N. Effects of gibberellic acid on water uptake and germination of sweet sorghum seeds under salinity stress. Chil. J. Agric. Res. 2019, 793, 415–424.

13. Finch-Savage WE, Bassel GW. Seed vigour and crop establishment: extending performance beyond adaptation. J. Exp. Bot. 2016; 673: 567–591.

14. Gadwal R, Naik GA. comparative study on the effect of salt stress on seed germination and early seedling growth of two Hibiscus species. 2014.

15. Botía P, Carvajal M, Cerdá A, Martínez V. Response of eight Cucumis melo cultivars to salinity during germination and early vegetative growth. Agron. Afr. 1998; 18: 503–513.

16. Gilliham M, Able JA, Roy SJ. Translating knowledge about abiotic stress tolerance to breeding programmes. The Plant. 2017; 90: 898–917.

17. Zhang H, Liu XL, Zhang RX, Yuan HY, Wang MM, Yang HY, Root damage under alkaline stress is associated with reactive oxygen species accumulation in rice (Oryza sativa L.). Front. Plant Sci. 2017; 8: 15–80.

18. Bose J, Munns R, Shabala S, Gilliham M, Pogson B, Tyerman SD, Chloroplast function and ion regulation in plants growing on saline soils: lessons from halophytes. J. Exp. Bot. 2017; 68: 3129–3143.

19. Shabala S, White RG, Djordjevic MA, Ruan YL, Mathesius U. Root-to-shoot signalling: integration of diverse molecules, pathways and functions. Funct. Plant Biol. 2015; 43: 87–104.

20. Jin X, Liu T, Xu J, Gao Z, Hu X. Exogenous GABA enhances muskmelon tolerance to salinity-alkalinity stress by regulating redox balance and chlorophyll biosynthesis. BMC plant biol. 2019; 19: 11–15.

21. Wu H, Zhang X, Giraldo JP, Shabala S. It is not all about sodium: revealing tissue specificity and signalling roles of potassium in plant responses to salt stress. Plant soil. 2018; 431: 1–17.

22. Lv BS, Li XW, Ma HY, Sun Y, Wei LX, Jiang CJ. Differences in growth and physiology of rice in response to different saline-alkaline stress factors. J. Agron. 2013; 105: 1119–1128.

23. Wang HL, Wang L, Tian CY, Huang ZY. Germination dimorphism in Suaeda acuminata: a new combination of dormancy types for heteromorphic seeds. S. Afr. J. Bot. 2012; 78: 270–275.

24. Kasote DM, Katyare SS, Hegde MV, Bae H. Significance of antioxidant potential of plants and its relevance to therapeutic applications. Int. J. Biol. Sci. 2015; 11: 9–82.

25. Mansour MMF, Ali EF. Evaluation of proline functions in saline conditions. Phytochem Rev. 2017; 140: 52–68.

26. Xu S, Hu B, He Z, Ma F, Feng J, Shen W, Yang J. Enhancement of salinity tolerance during rice seed germination by presoaking with hemoglobin. Int. J. Mol. Sci. 2011; 124: 2488–2501.

27. Akbarimoghaddam H, Galavi M, Ghanbari A, Panjehkeh N. Salinity effects on seed germination and seedling growth of bread wheat cultivars. Trakia J. Sci. 2011; 19: 43–50.

28. Carpýcý E, Celyk N, Bayram G. Effects of salt stress on germination of some maize (Zea mays L.) cultivars. Afr. J. Biotechnol. 2009; 8–19.

29. Khodarahmpour Z, Ifar M, Motamedi M. Effects of NaCl salinity on maize (Zea mays L.) at germination and early seedling stage. Afr. J. Biotechnol. 2012; 112: 298–304.

30. Akram N A, Jamil A. Appraisal of physiological and biochemical selection criteria for evaluation of salt tolerance in canola (Brassica napus L.). Pak. J. Bot. 2007; 395: 1593–1608.

31. Abid M, Malik SA, Bilal K, Wajid RA. Response of Okra (Abelmoschus esculentus L.) to EC and SAR of Irrigation Water. Int. J. Agric. Biol. 2002; 43: 311–314.

32. Aaron TA, Elvis AB, Faustina A, Kingsley T, Edmund O. Phenotypic traits detect genetic variability in Okra (Abelmoschus esculentus. L. Moench). Afr. J. Agric. Res. 2016; 33: 3169–3177.

33. Esan AM, Masisi K, Dada FA, Olaiya CO. Comparative effects of indole acetic acid and salicylic acid on oxidative stress marker and antioxidant potential of okra (Abelmoschus esculentus L) fruit under salinity stress. Sci. Horti. 2017; 216: 278–283.

34. Kader MA, Lindberg S. Uptake of sodium in protoplasts of salt-sensitive and salt-tolerant cultivars of rice, (Oryza sativa L). determined by the fluorescent dye SBFI. J. Exp. Bot. 2005; 422: 3149–3158.

35. Kumar R, Shamet GS, Mehta H, Alam NM, Tomar JM, Chaturvedi OP, Khajuria N. Influence of gibberellic acid and temperature on seed germination in Chilgoza pine (Pinus gerardiana Wall.). Indian J. Plant Physiol. 2014; 194: 363–367.

36. Dubois M, Gilles KA, Hamilton JK, Rebers PT, Smith F. Colorimetric method for determination of sugars and related substances. Anal. chem. 1956; 28: 350–356.

37. Bates LS, Waldren RP, Teare I. Rapid determination of free proline for water-stress studies. Plant soil. 1973; 39: 205–207.

38. Arnon DI. Copper enzymes in isolated chloroplasts. Polyphenoloxidase in Beta vulgaris. Plant physiol. 1949; 24: 1.

39. Shin YK, Bhandari SR, Jo JS, Song JW, Cho MC, Yang EY, Lee JG. Response to salt stress in lettuce: Changes in chlorophyll fluorescence parameters, phytochemical contents, and antioxidant activities. Agronomy. 2020; 10: 16–27.

40. Tarchoun N, Saadaoui W, Mezghani N, Pavli OI, Falleh H, Etropoulos SA. The Effects of Salt Stress on Germination, Seedling Growth and Biochemical Responses of Tunisian Squash (Cucurbita maxima Duchesne) Germplasm. Plants, 2022; 11: 800.

41. Srivastava S, Shrama P. Effect of NaCl on chlorophyll fluorescence and thylakoid membrane proteins in leaves of salt sensitive and tolerant rice (Ory;za sativa L) varieties. J. stress physiol. biochem. 2021; 17: 35–44.

42. Janghel DK, Kumar K, Sunil R, Chhabra AK. Genetic diversity analysis, characterization and evaluation of elite chickpea (Cicer arietinum L.) genotypes. Int. J. Curr. Microbiol. App. Sci. 2020; 90: 199–209.

43. Hasan B, Higano Y, Yabar H, Devkota M, Lamers J. Conservation agriculture practices in salt-affected, irrigated areas of Central Asia: Crop price and input cost variability effect on revenue risks. Sustain. Agric. Res. 2015; 526: 2016–37925.

44. Bayuelo-Jiménez JS, Craig R, Lynch JP. Salinity tolerance of Phaseolus species during germination and early seedling growth. Crop Sci. 2002; 42: 1584–1594.

45. Cokkizgin A. Salinity stress in common bean (Phaseolus vulgaris L.) seed germination. Not. Bot. Horti. Agrobot. Cluj. Napoca. 2012; 40: 177–182.

46. Meot-Duros L, Magne C. Effect of salinity and chemical factors on seed germination in the halophyte *(Crithmum maritimum* L.) Plant Soil. 2008; 313: 83–87.

47. Jamil M, Rha ES. The effect of salinity (NaCl) on the germination and seedling of sugar beet *(Beta vulgaris* L.) and cabbage *(Brassica oleracea* L.). Plant Genet. Resour. 2004; 73: 226–232.

48. Foti C, Khah E, Pavli O. Germination profiling of lentil genotypes subjected to salinity stress. Plant Biol. 2019; 21: 480–486.

49. Jamil M, Lee CC, Rehman SU, Lee DB, Ashraf M, Rha E.S. Salinity (NaCl) tolerance of Brassica species at germination and early seedling growth. Elec. J. Env. Agricult. Food Chem. 2005, 44, 970–976.

50. Munns R. Comparative physiology of salt and water stress. Plant cell environ. 2002; 25: 239–250.

51. Gama PBS, Inanaga S, Tanaka K, Nakazawa R. Physiological response of common bean (Phaseolus vulgaris L.) seedlings to salinity stress. Afr. J. biotechnol. 2007; 6–2.

52. Ndakidemi PA, Makoi JHJR. Effect of NaCl on the productivity of four selected common bean cultivars (Phaseolus vulgaris L.). 2009;4: 1066–1072.

53. Szabados L, Savouré A. Proline: a multifunctional amino acid. Trends in plant sci. 2010 15: 89–97.

54. Kavi-kishor PB, Sreenivasulu N. Is proline accumulation per se correlated with stress tolerance or is proline homeostasis a more critical issue? Plant cell environ. 2014; 37: 300–311.

55. Akram NA, Iqbal M, Muhammad A, Ashraf M, Al-Qurainy F, Shafiq S. Aminolevulinic acid and nitric oxide regulate oxidative defense and secondary metabolisms in canola (Brassica napus L.) under drought stress. Protoplasma. 2018; 255: 163–174.

56. Iqbal N, Ashraf MY, Javed F, Martinez V, Ahmad K. Nitrate reduction and nutrient accumulation in wheat grown in soil salinized with four different salts. J. plant nutr. 2006; 29: 409–421.

57. Nekir B, Mamo L, Worku A, Bekele T. Evaluation of wheat varieties/lines for salt tolerance at different growth stages. Greener J. Soil Sci. Plant Nutr. 2019; 61: 1–7.

58. Kaouther Z, Mariem BF, Fardaous M, Cherif H. Impact of salt stress (NaCl) on growth, chlorophyll content and fluorescence of Tunisian cultivars of chili pepper (Capsicum frutescens L.). J. stress physiol. biochem. 2012; 84: 236–252.

